# ViralMap: Predicting Features in Viral Proteins from Primary Sequence

**DOI:** 10.64898/2026.04.07.716565

**Authors:** Shrish Dwivedi, Shaunak Kar, Andrew P. Horton, Jimmy D. Gollihar

## Abstract

Modern viral vaccines are designed to elicit an immune response against viral proteins that mediate infection, making those proteins important targets for characterization and engineering. To improve vaccine efficacy, the proteins often require changes to specific residues or domains to improve immunogenicity and induce a protective response. These engineering strategies vary significantly across viruses, and comprehensive and accurate protein sequence annotation is crucial for guiding vaccine design. The growing risk of novel pathogen emergence and initiatives such as the CEPI 100 Days Mission to rapidly counter “Disease X” threats heighten the need for tools that can convert viral protein sequences from newly characterized genomes or emerging variants into the annotation profiles required for antigen engineering. To address this, we developed ViralMap, a multi-label annotation model tailored for eukaryotic viral proteins. By leveraging ESM-2 language model representations, ViralMap simultaneously predicts ten distinct annotation classes spanning domain topology and localization, post-translational modifications, and structural features directly from primary sequences. The model achieves a residue-level precision-recall area under the curve (PR-AUC) of 0.75 or greater for seven of the ten classes and realizes predictive performance competitive with established tools across the eight benchmarked classes. Case studies on complex glycoproteins, including the SARS-CoV-2 spike and HIV-1 Env, illustrated the model’s ability to generalize across viral strains and to novel viral families not seen during training. By providing a unified, sequence-based framework for multi-label annotation, ViralMap offers a practical and scalable bridge from raw viral protein sequences to the annotation profiles required for antigen engineering.

## INTRODUCTION

Vaccines function by exposing the body to pathogen signatures, or antigens, to elicit a protective immune response against future disease. For viruses, the most relevant antigens are proteins that play a role in infection, and these are prime targets for vaccine engineering.^1^ For example, the severe acute respiratory syndrome coronavirus 2 (SARS-CoV-2) spike glycoprotein, which mediates viral entry by binding to host-cell receptors,^2^ was the major component of all five vaccines recommended by the World Health Organization in 2022. Translating a viral protein into an effective vaccine often requires modifying the protein sequence to improve its conformational stability, epitope presentation and expression, and ultimately to induce a robust and protective immune response.^3^ Such design strategies have included stabilizing the extracellular domains of the SARS-CoV-2 spike protein^4^ and the respiratory syncytial virus (RSV) fusion protein in pre-fusion conformations,^5^ as well as removing glycosylation sites on the human immunodeficiency virus type-1 (HIV-1) envelope protein to expose antibody targets.^6,7^ Establishing a comprehensive and accurate sequence annotation profile for viral proteins is therefore imperative for understanding viral function and supporting vaccine development.

An annotation profile encompasses a diverse set of residue- and domain-level features. Transmembrane domains anchor proteins in membranes and define the boundaries between cytoplasmic and extracellular regions. These topological features are central to antigen design strategies such as extracellular domain stabilization and transmembrane deletion.^3^ Signal peptides are primarily N-terminal regions that direct proteins to the secretory pathway,^8^ and their modification has been shown to improve immunogenicity.^9^ Before reaching their functional form, many viral surface proteins undergo proteolytic maturation in which a precursor polypeptide is cleaved into distinct subunits by host proteases such as furin.^2,10,11^ Post-translational modifications further shape antigen properties: N-glycosylation influences protein folding, receptor binding, and immune evasion;^12,13^ because glycans can shield or expose epitopes, glycosylation site modification is a key lever in antigen design.^6,7^ Disulfide bonds stabilize protein tertiary structure and are similarly exploited in antigen design to lock proteins in preferred conformations.^3^ Additional structural features, including coiled coils in viral fusion machinery^14,15^ and intrinsically disordered regions prevalent in viral proteomes,^16^ further inform stabilization and design strategies for viral antigens. Characterizing these features across a viral protein provides a foundation for downstream engineering decisions.

The need to generate this functional profile rapidly is heightened by the growing risk of infectious disease outbreaks largely driven by anthropogenic factors.^17^ This risk has motivated significant investment in global pandemic preparedness initiatives, such as the Coalition for Epidemic Preparedness Innovations (CEPI) 100 Days Mission to counter novel (“Disease X”) pathogens.^18^ The initiative aims to develop capabilities to create and manufacture a vaccine against any unspecified pathogen within 100 days of its recognition, and retrospective studies have concluded that achieving this timeline during the SARS-CoV-2 pandemic would have saved ∼8 million lives.^19^ Advances in metagenomic sequencing^20^ and bioinformatics^21^ have streamlined the recovery of putative protein sequences from newly characterized viral genomes. A critical bottleneck is therefore the conversion of extracted viral protein sequences into actionable targets for antigen engineering through functional annotation.

Machine learning offers an effective alternative to labor-intensive experimental characterization and traditional alignment-based methods for protein sequence annotation. Several models have been developed for specific annotation tasks, such as predicting signal peptides,^22,23^ intrinsically disordered regions,^24,25^ and N-glycosylation sites.^26,27^ Some approaches also predict multiple annotation types simultaneously, particularly for topology-related features such as transmembrane, cytoplasmic and extracellular domain localization.^28,29^ However, these models must be combined to obtain more comprehensive annotation profiles, and constructing such multi-tool pipelines is technically demanding due to incompatible dependencies, formats, software availability and licensing restrictions. Furthermore, most general protein annotation tools are developed without explicit specialization for viral sequences, whose rapid evolution drives substantial sequence divergence from host proteins,^30^ and can limit the generalizability of models trained predominantly on non-viral datasets^31^, which reduces the utility of alignment-based approaches^32^ when homologous relationships are weak or absent. Existing virus-specific annotation models are primarily focused on bacteriophages,^33–36^ which differ in key biological properties from the eukaryotic viruses most relevant to human health and vaccine development. For instance, phage virion glycosylation is rare,^37^ whereas N-linked glycosylation is pervasive among many eukaryotic viruses and plays important roles in host-virus interactions.

These limitations underscore the need for a viral annotation model that bridges the critical gap between genomic characterization and antigen engineering. Specifically, it requires a model that (1) simultaneously predicts multiple annotation classes relevant to antigen engineering, (2) is tuned to eukaryotic viral protein sequences, and (3) achieves performance competitive with established specialized tools when used for the same objective. To this end, we developed ViralMap, an annotation model for eukaryotic viral proteins that predicts, at residue-level resolution, ten annotation classes spanning three functional categories: topology and localization (transmembrane, cytoplasmic, and extracellular domains), post-translational modifications (N-glycosylation, furin cleavage, chain cleavage, and disulfide bond sites), and structural features (coiled coils and intrinsically disordered regions). By formulating annotation as a multi-label residue classification task, ViralMap captures overlapping, biologically coherent annotations within a single model. Further, because ViralMap requires only primary protein sequences to generate these predictions, it enables immediate functional profiling of novel viruses from newly extracted genomic data.

We show that ViralMap achieves competitive performance across these diverse annotation classes when evaluated alongside a suite of established tools on eukaryotic viral proteins. By replacing fragmented multi-tool pipelines with a single, sequence-based framework, ViralMap provides a practical and scalable bridge from raw viral protein sequences to the annotation profiles that inform antigen engineering efforts.

## METHODS

### Data selection

All eukaryotic (non-phage) viral proteins were downloaded from UniProt using taxonomy ID 10239, excluding organisms containing the keyword ‘phage’ (n=4,090,737 proteins; 15,488 Swiss-Prot). Annotation quality in this initial set was generally poor, necessitating a customized curation pipeline. Proteins were first filtered by annotation score (≥4.0) and the presence of a non-null ‘Features’ field; all Swiss-Prot entries were retained regardless of these criteria. Keyword and length filters (100–1,024 residues; upper bound set by the ESM-2 context window) removed residual phage entries, fragments, and long polyproteins, yielding 300,406 proteins. These were clustered with MMseqs2 single-linkage clustering (60% sequence identity, 50% bidirectional coverage, E-value ≤0.001), and an annotation-aware selection algorithm **(Supplementary Note 1)** was applied within each cluster to select for cluster representatives. The final dataset comprised 8,238 proteins totaling 3,131,029 residues **(Supplementary Table 2)**.

### Annotation class representation

All annotation classes were converted into binary labels using the coordinates provided in UniProt. Each protein sequence of length T was therefore associated with a binary label tensor *Y* ∈ {0,1}^Tx10^, where *Y_i,k_* = 1 indicates that residue *i* is positively annotated for annotation class *k.* Except for chain cleavage sites and topology, all annotation coordinates were used as provided by UniProt. For chain cleavage sites, UniProt provides (start, end) positions to indicate mature polypeptide chains. To predict the boundaries between these regions, we inverted the provided coordinates so that the paired residues marking the end of one chain and the start of the next were labeled as 1 in the tensor. Because chain annotations denote the boundaries of mature polypeptides regardless of the specific protease mechanism, these sites naturally encompass and overlap with explicitly annotated protease targets, such as furin cleavage sites. For topology, all residues annotated as “Cytoplasmic” or “Intravirion” were mapped to the cytoplasmic class, and residues annotated as “Extracellular”, “Virion surface”, or “Lumenal” were mapped to the extracellular class. Rare instances of topology corresponding to “Nuclear” or “Perinuclear space” were grouped with the cytoplasmic class for consistency. All transmembrane domains in the final dataset were alpha-helical; beta-barrel transmembrane domains are outside the current scope. Isolated instances of different N-glycosylation subtypes (hybrid, complex, and high mannose) were mapped to the N-glycosylation class.

### Data splitting

The final dataset was divided into five folds for cross-validation. Folds were assigned in a viral-family-aware manner to help balance taxonomic bias, while also balancing residue-level counts of all annotation classes across folds **(Supplementary Table 2)**. Large viral families with more than 200 proteins were distributed across multiple folds, while smaller families were grouped into individual folds **(Supplementary Fig. 6)**. In either case, the clusters created during representative selection were assigned entirely to a single fold to prevent homology leakage. This splitting strategy ensured that the overall test performance reflected both cross-strain generalization (as homolog clusters within large families were kept strictly separated) and novel-family generalization (as smaller families were isolated entirely). For each cross-validation iteration during training, three folds were used for training, one for validation (early stopping), and one for test evaluation. For testing, we used MMseqs2 easy-search to compare test-fold sequences with their corresponding training and validation sequences to ensure strict homology separation. For each test fold, isolated proteins that exceeded the identity threshold with any training or validation sequence were removed prior to evaluation. This had minimal impact on the test set sizes (n=5 to 29 proteins removed depending on the test fold).

### Model architecture

ViralMap is a two-stage architecture **(Fig. 1b)** comprising a base model and a post-processing module. The base model consists of the pretrained 33-layer, 650-million-parameter ESM-2 protein language model with a neural network classification head. To preserve general learned representations, all but the final four transformer layers are frozen. For a given input sequence, the language model outputs 1,280-dimensional embeddings for each residue, which are then passed to a 2-layer fully connected neural network with dropout (p=0.3) and ReLU activation. The final output of the base model is 10 probabilities per residue position after sigmoid activation. Because the base model is trained using a binary cross-entropy objective, the outputs represent independent class probabilities at each residue position.

**Figure 1.**
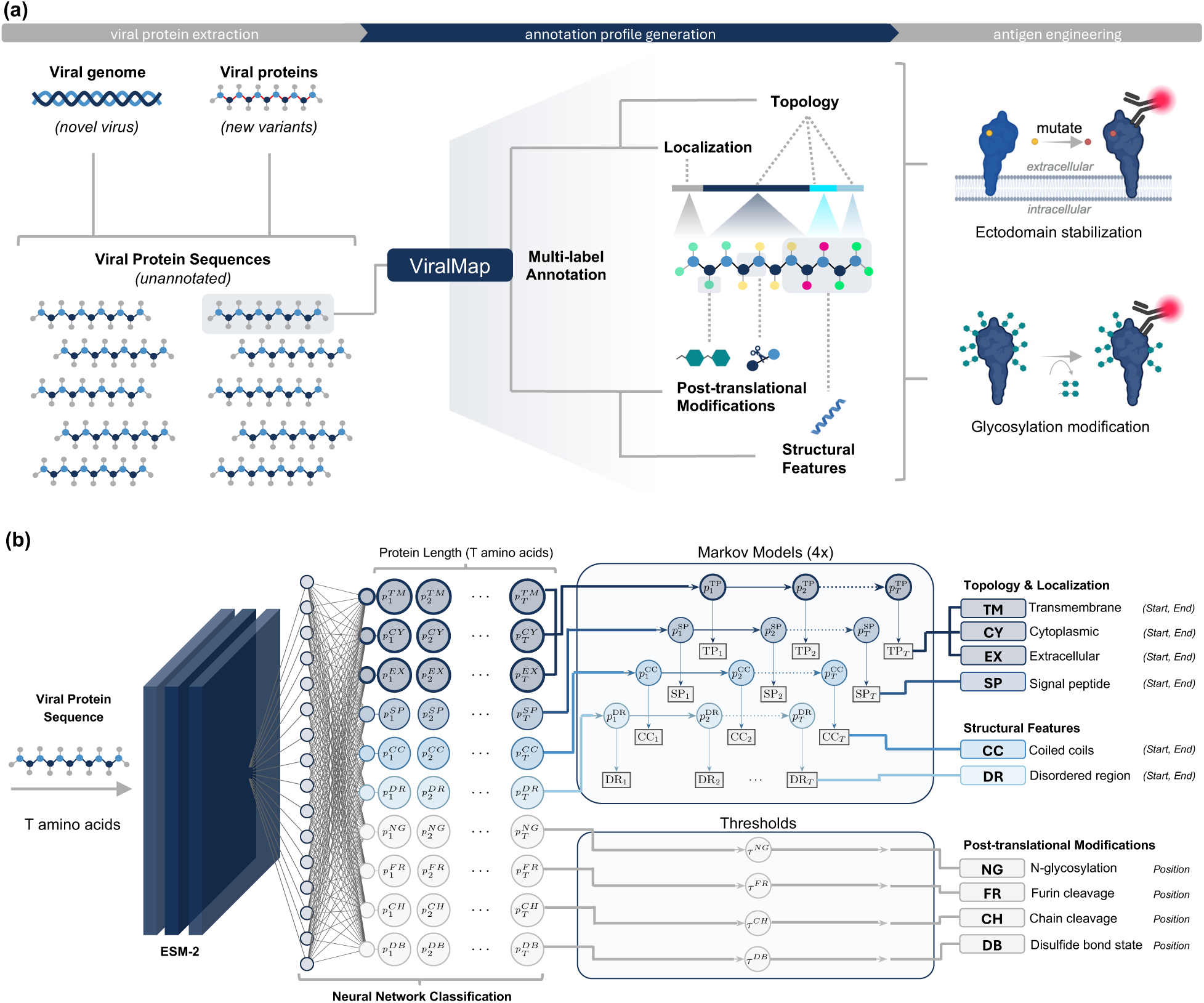
ViralMap overview and architecture. **(a)** End-to-end workflow: unannotated viral protein sequences are processed by ViralMap to produce annotation profiles spanning ten classes across three categories that support antigen engineering decisions. **(b)** Model architecture: the base model (ESM-2 with a neural-network classification head) produces per-residue probabilities for all ten annotation classes, which are post-processed into (start, end) region coordinates by HMMs or individual residue positions using probability thresholds (*τ*).

The post-processing module produces binary predictions from the base model probabilities for each residue and class. For individual residue annotation classes (N-glycosylation, furin cleavage sites, disulfide bond cysteines, and chain boundaries), an F2-optimized threshold calculated on the validation set is applied. Although UniProt annotates furin cleavage sites using two adjacent residues, we treat the class as a single residue prediction. For annotation classes requiring start and end positions (cytoplasmic domains, extracellular domains, transmembrane domains, signal peptides, coiled coils, and disordered regions), the base model probabilities are decoded using Hidden Markov Models (HMMs) with the Viterbi algorithm.

Four HMMs were implemented: a 4-state topology model over {None, Cytoplasmic, Extracellular, Transmembrane} that enforces biological mutual exclusivity, and three 2-state models: {None, Signal peptide}, {None, Coiled coil}, and {None, Disordered}. For each fold, transition matrices and start priors were calculated from the training data. For each HMM, base model probabilities were converted to log-emissions:

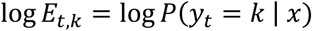

where *P*(*y_t_* = *k* | *x*) is the base model’s predicted probability for class *k* at position *t* in the protein sequence, and P(None) = 1 − ∑*_k_P*(*y_t_* = *k*). The None state in each HMM represents residues without the corresponding annotation. To improve sensitivity, the None-state log-emissions were penalized by 4.0 for the topology HMM and 1.0 for the two-state HMMs. Contiguous runs of non-None states in the decoded sequence define predicted annotation regions with explicit start and end positions. In this iteration of ViralMap, the HMMs function solely as structured decoders that enforce specific biological constraints, generate start and end coordinates, and preserve the base model’s underlying discriminative performance.

### Training

The base model was trained on all classes simultaneously at the residue level, using binary cross-entropy loss with weighting schemes to address both inter- and intra-class imbalance. Inter-class imbalance refers to the positive residue skew between classes, while intra-class imbalance refers to the overwhelming negative residues compared to positive within each class. To address intra-class imbalance, positive residues were upweighted using:

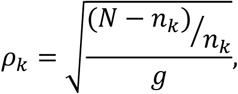

where *N* is the total number of residues, *g* is the geometric mean of the raw positive-negative ratios across classes, and *n_k_* is the number of positive residues for class *k*. For inter-class imbalance, class weights were computed using the effective number of samples framework:^38^

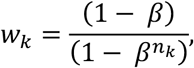

with *β* = 1 – 10^−6^. The binary cross-entropy loss with sigmoid (*σ*) activation with class-weighting was therefore:

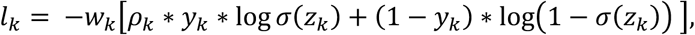

where *z_k_* is the logit output for class *k* from the base model. All class weights were computed using only the training folds for a given cross-validation run. The base model was trained using AdamW^39^ optimization with a learning rate 3*x*10^−4^ and 10% linear warmup. Training used mixed precision (fp16) with gradient clipping (max norm=0.5). Early stopping with patience of 4 was determined by the macro-averaged PR-AUC on the held-out validation folds. All training was performed on Nvidia GH200 GPUs using PyTorch 2.5.1.

### Evaluation

ViralMap was evaluated against established annotation tools for all classes except chain cleavage sites and disulfide bond states, for which no comparable sequence-based tools (requiring only sequences, not structures) were available. Benchmark tools were selected based on three criteria: (1) publication or preprint available, (2) clear documentation, and (3) command-line interface capable of processing the full ViralMap dataset (8,238 sequences). Several recently published tools were excluded primarily due to unclear documentation or a lack of scalable interfaces. The final comparison tools in the benchmark were: DeepTMHMM for signal peptides, transmembrane domains, cytoplasmic domains, and extracellular domains;^29^ DeepCoil for coiled coils;^40^ AIUPred for disordered regions;^41^ NetNGlyc for N-glycosylation sites;^42^ and ProP for furin cleavage sites.^43^ All tools were run with default parameters and converted into binary predictions as described in **Supplementary Table 3.**

We evaluated performance at the residue level. Precision and recall are computed directly from per-residue predictions without boundary adjustments. The formulae for precision and recall are as follows:

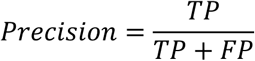

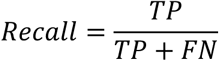

True positives (TP) are residues correctly predicted as positive that match UniProt annotations. False positives (FP) are residues predicted as positive that lack corresponding UniProt annotations. False negatives (FN) are UniProt-annotated residues that the models fail to predict. Because UniProt annotations are incomplete, apparent FP predictions often include correct predictions for annotations absent from the database, a limitation that affects all supervised methods evaluated against curated datasets. We therefore interpret the reported precision metrics as a loose lower bound of true precision.

Precision-recall area under the curve (PR-AUC) is reported for ViralMap only, as benchmark tools do not uniformly provide per-residue probabilities. For ViralMap post-translational modification classes, F2-optimized thresholds (prioritizing recall) were applied **(Table 1)**. We evaluated strictly at the residue level because it implicitly captures protein-level count accuracy and boundary fidelity. For topology classes (transmembrane, cytoplasmic, extracellular), residue-level metrics were computed only on residues with at least one topology annotation, as unlabeled residues in such proteins most likely represent missing annotations rather than true negatives. All metrics are reported as mean ± standard deviation across the five test folds.

**Table 1.**
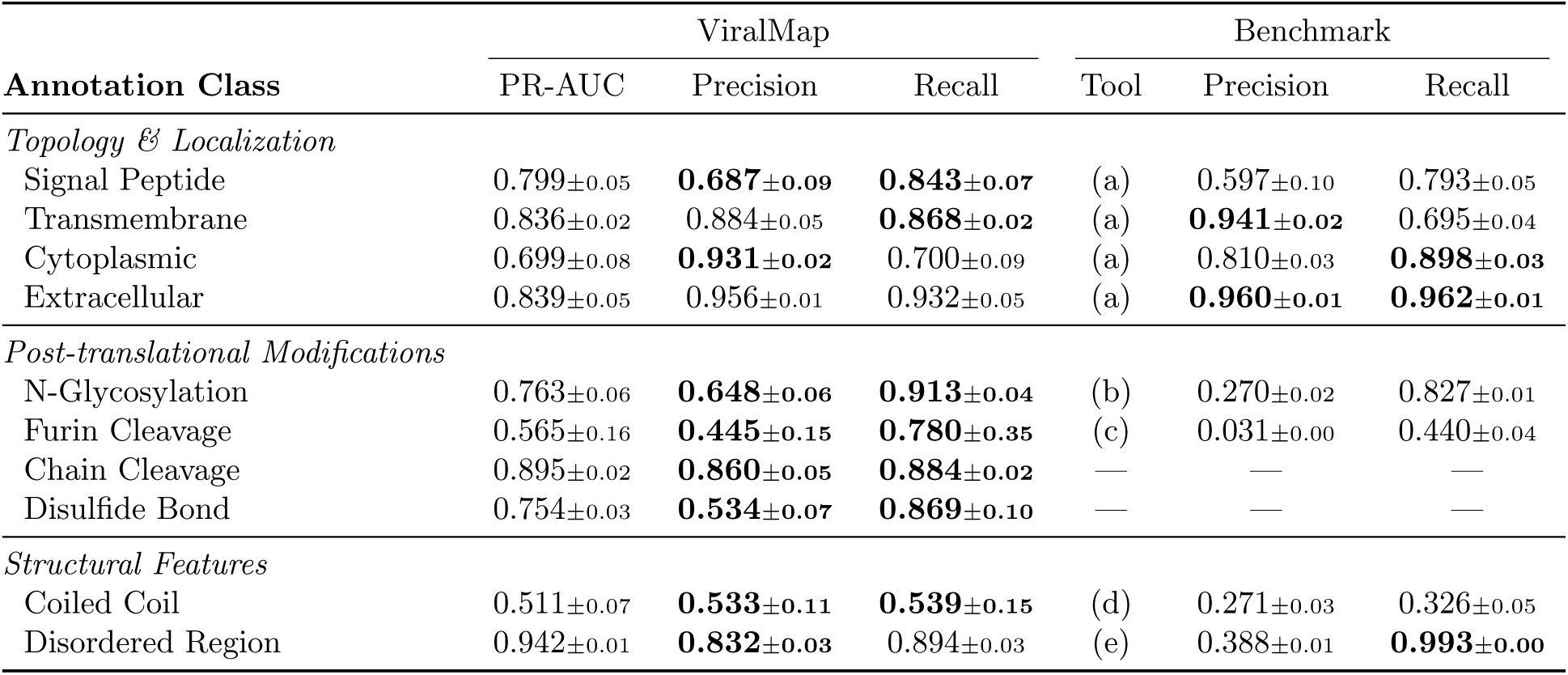
Residue-level classification performance of ViralMap and benchmark tools on held-out eukaryotic viral proteins. Benchmark tools: **(a)** DeepTMHMM, **(b)** NetNGlyc, **(c)** ProP, **(d)** DeepCoil, **(e)** AIUPred. All metrics are reported as mean ± standard deviation across five test folds. PR-AUC is reported for the ViralMap base model only; precision and recall are reported after post-processing (HMMs for topology & localization and structural features; F2-optimized thresholds for post-translational modifications. **Supplementary Table 8** reports ViralMap performance for post-translational modifications using a standard 0.5 threshold.) Topology metrics were computed only on proteins with at least one topology annotation in UniProt. Bold indicates the higher value between ViralMap and the benchmark tool.

## RESULTS

### ViralMap overview

ViralMap is designed to convert unannotated viral protein sequences into residue-level annotation profiles that can inform downstream antigen engineering **(Fig. 1a)**. In practice, these sequences may originate from a newly characterized viral genome or from new variants of a known virus. In both instances, the input to ViralMap is the primary protein sequence alone, and the output is a multi-label annotation profile spanning ten classes across three categories **(Fig. 1b)**. For topology and localization, the model predicts transmembrane, cytoplasmic, and extracellular domain boundaries as well as signal peptides. For post-translational modifications, it identifies N-glycosylation sites, furin cleavage sites, and disulfide bond sites, where the model predicts whether individual cysteines participate in disulfide bonds rather than predicting explicit bond pairings between cysteine residues.^44^ The model also predicts chain cleavage sites, which represent boundaries between mature polypeptides regardless of the specific protease mechanism, providing broader coverage of viral protein maturation events. Finally, for structural features, the model localizes coiled coil domains and intrinsically disordered regions. By producing all ten annotations simultaneously from sequence alone, ViralMap provides a unified framework for generating the annotation profiles needed to support antigen engineering decisions.

The underlying ViralMap architecture **(Fig. 1b)** is a two-stage pipeline. The base model pairs the pretrained 33-layer, 650-million-parameter ESM-2^45^ protein language model (PLM) with a neural-network classification head, producing per-residue probability estimates for each annotation class. PLMs such as ESM-2 learn contextualized representations of protein sequences and have been successfully applied to viral proteins for tasks such as predicting antigen fitness^46^ and immune escape.^47^ The per-residue probabilities are then post-processed to produce concrete predictions while enforcing biological constraints such as mutual exclusivity among topological domains. Classes requiring start and end positions (topology, localization, and structural features) are decoded using Hidden Markov Models (HMMs), while individual-residue classes (post-translational modifications) are converted into binary predictions using probability thresholds. Full architectural and training details are described in the methods. ViralMap is implemented in Python and uses a command-line interface.

### Annotation-aware dataset curation

Training ViralMap involved balancing multiple conflicting goals during dataset curation: minimizing false negatives caused by missing or incomplete annotations, prioritizing proteins with higher-quality evidence in their annotations, retaining isolated examples of rare annotation classes, and reducing sequence redundancy while preserving enough related proteins to use natural sequence variation as a form of data augmentation. Because ViralMap predicts ten annotation classes simultaneously, dataset curation could not rely solely on sequence-based redundancy reduction; protein selection also needed to account for annotation quality and coverage across classes. We downloaded all eukaryotic (non-phage) viral proteins from UniProt^48^ (n=4,090,737) and found that annotation quality was generally poor: over 90% (n=3,686,285 proteins) had annotation scores of ≤3.0, where the best-annotated entries in UniProt have a score of 5.0 on a 1-5 scale **(Supplementary Fig. 1)**. Including poorly annotated proteins would inherently introduce noise since missing annotations would be incorrectly interpreted by the model as true negatives during training. Simply restricting the dataset to Swiss-Prot (reviewed) entries would not address the false-negative problem since over 98% of the highest-scoring entries (annotation score 5.0; n=128,901 proteins) were TrEMBL (unreviewed) records **(Supplementary Fig. 1)**. These limitations necessitated a customized dataset curation pipeline **(Fig. 2)** that captured annotation-level presence and quality across all classes of interest simultaneously.

**Figure 2.**
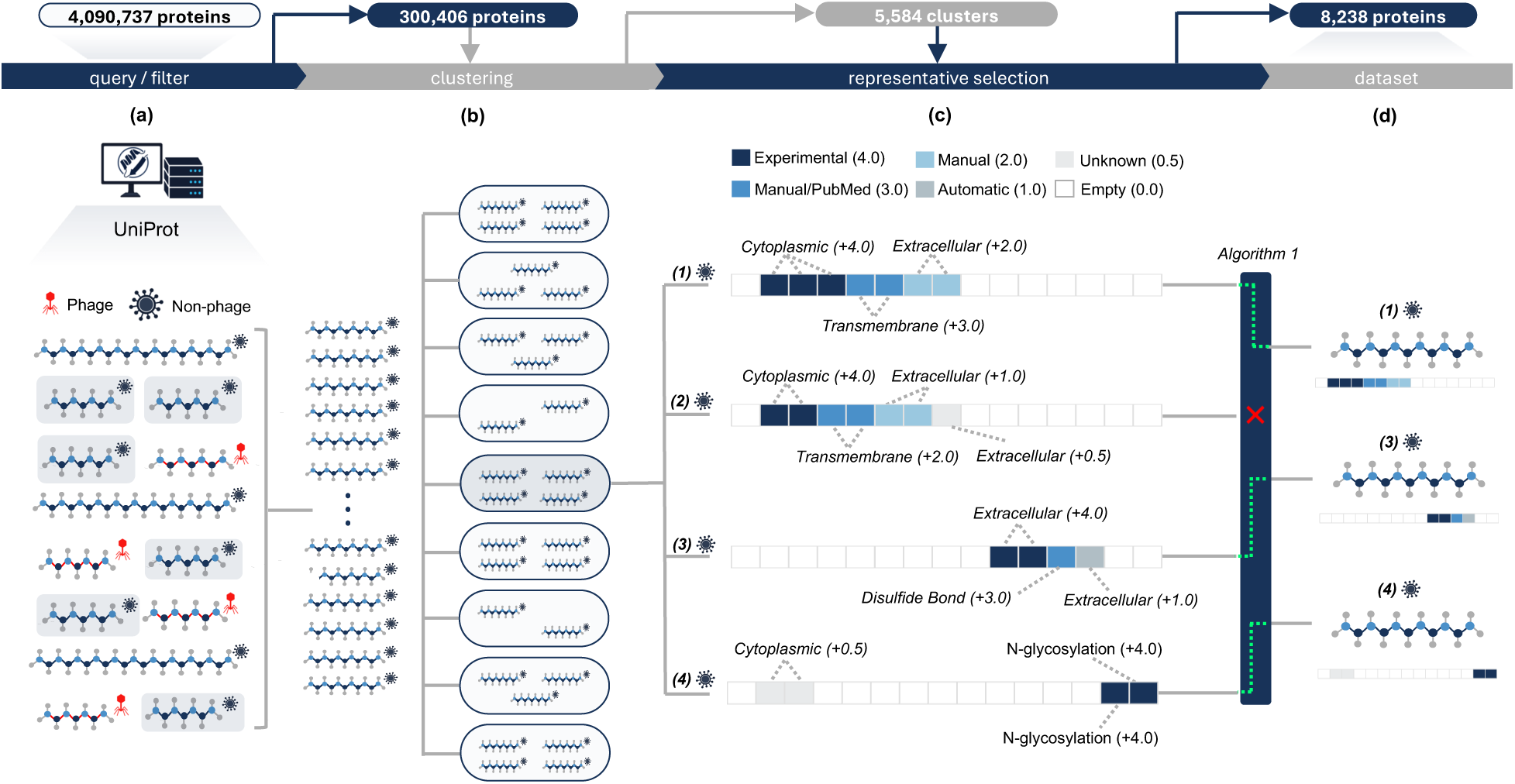
Preprocessing and annotation-aware dataset curation for ViralMap. **(a)** Eukaryotic viral proteins were retrieved from UniProt and filtered by annotation quality, sequence length, and taxonomy to exclude residual phage records. **(b)** Filtered proteins were clustered by sequence identity. **(c)** A representative selection algorithm **(Algorithm 1, Supplementary Note 1)** selected candidates based on annotation evidence quality and spatial novelty to reduce redundancy while preserving overall annotation coverage within the cluster, as illustrated here for a hypothetical cluster of four proteins. **(d)** The final reduced-redundancy dataset preserves complementary residue-level annotations for downstream model training.

The pipeline proceeded in four stages **(Fig. 2a–d)**. Initial filtering **(Fig. 2a)** retained proteins with annotation scores of 4.0 or higher and a non-null ‘Features’ field, which UniProt uses to summarize annotation counts across classes for a given protein. Additional keyword filtering removed residual phage entries, and a length filter retained proteins between 100 and 1,024 residues to exclude fragments and excessively long polyproteins. This yielded 300,406 proteins. To reduce redundancy, these proteins were then clustered **(Fig. 2b)** using MMseqs2^49^ single-linkage clustering at 60% identity with 50% bidirectional alignment coverage (E-value cutoff of 0.001). Clustering produced largely homogeneous groups with respect to protein type and viral identity **(Supplementary Figs. 2 and 3**).

**Figure 3.**
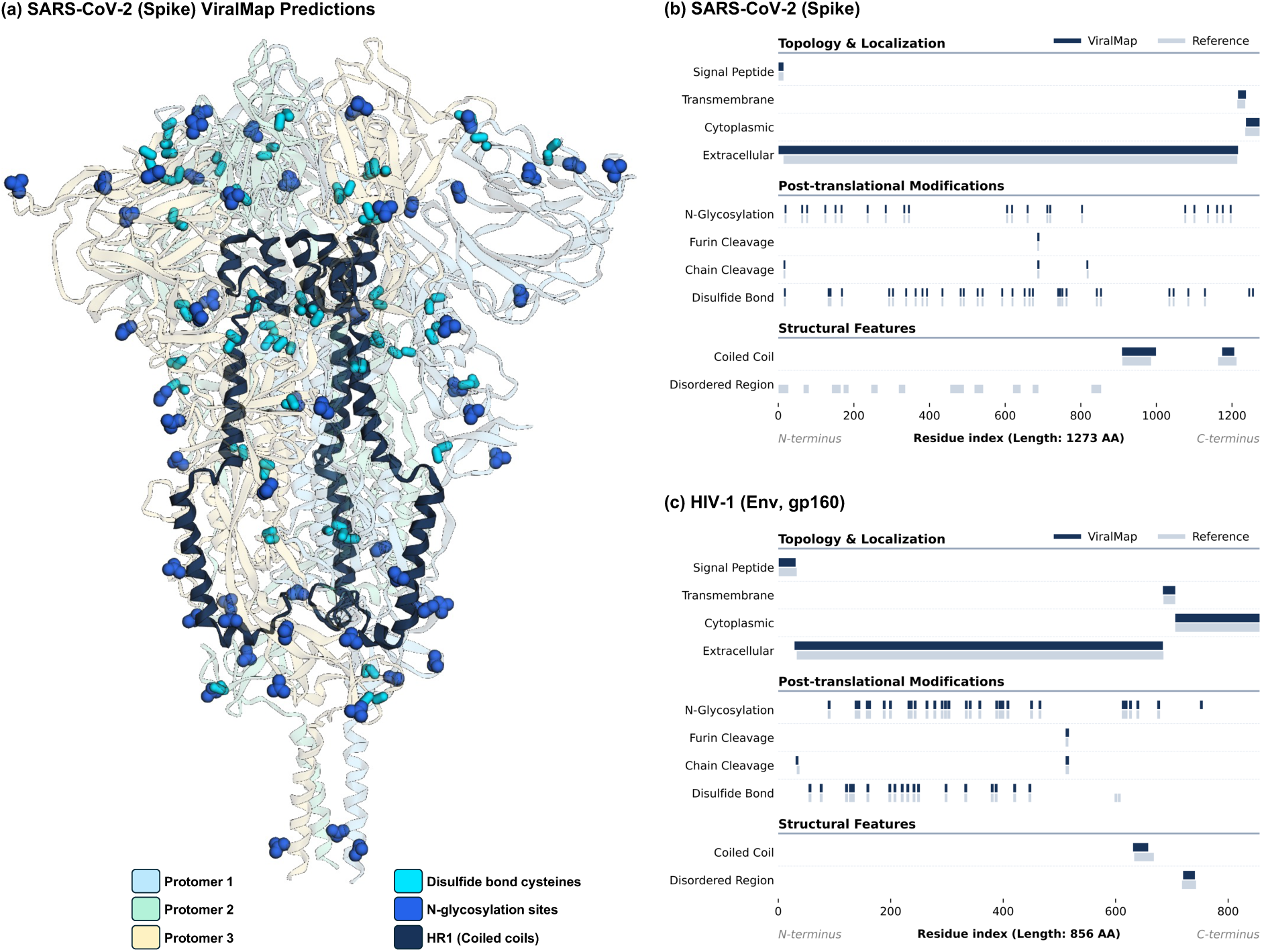
Visualization of ViralMap predictions against reference annotations for key viral glycoproteins. **(a)** 3D spatial mapping of select ViralMap sequence annotations onto the prefusion structure of the SARS-CoV-2 spike glycoprotein trimer (PDB: 6XR8). **(b)** 1D sequence tracks comparing ViralMap predictions (dark bars) against reference annotations (light bars) for the SARS-CoV-2 spike protein (UniProt ID: P0DTC2). All reference annotations are sourced from UniProt, with the exception of the coiled-coil regions: heptad repeat 1 (HR1) and heptad repeat 2 (HR2) are derived from the cryo-EM structure of the SARS-CoV-2 spike protein.^51^ **(c)** 1D sequence tracks comparing ViralMap predictions and reference annotations for the HIV-1 envelope (gp160) glycoprotein (UniProt ID: P04578). All reference annotations are sourced from UniProt.

We then applied a custom representative selection algorithm within each cluster **(Fig. 2c; Supplementary Note 1)** designed to prioritize proteins with high-quality annotations while retaining complementary coverage across classes. Each protein was encoded as a fixed-length annotation vector, in which bin values reflected both the positions and the evidence quality of its annotations. To distinguish between annotation sources, the evidence quality of each individual annotation was scored using a five-point scale derived from Evidence and Conclusion Ontology (ECO)^50^ codes: annotations with experimental evidence received a score of 4, manual annotations with publication support received 3, manual annotations without publication support received 2, automatic annotations received 1, and annotations without ECO codes received 0.5 **(Supplementary Table 1)**. Representative proteins were then iteratively selected using an information-gain criterion that combined the annotation vector magnitude (overall quality and density) and the cosine distance from already-selected representatives (complementary coverage). Selection continued until a per-cluster cap was reached, or no remaining protein exceeded a minimum information-gain threshold.

The algorithm reduced the dataset from 300,406 to 8,238 proteins (3,131,029 residues; **Fig. 2d**) while preferentially retaining proteins with experimentally or manually validated annotations **(Supplementary Figs. 4 and 5)**. By allowing the selection of multiple representatives from the same cluster, this approach also preserved within-cluster sequence divergence as implicit data augmentation. The annotation instance and residue counts for the final dataset, stratified by evidence source and cross-validation fold, are reported in **Supplementary Table 2.**

### Classification performance

**Table 1** summarizes the residue-level classification performance across all annotation classes and compares performance for each class against established tools DeepTMHMM,^29^ DeepCoil,^40^ AIUPred,^41^ NetNGly,^42^ and ProP,^43^ where applicable. All classification metrics were computed on held-out test folds, with homology separation among the training, validation, and test sets. ViralMap achieved PR-AUC ≥ 0.75 for 7 of the 10 classes, indicating successful multi-label prediction despite severe class imbalance. Because UniProt annotations are incomplete, apparent false positives may include correct predictions for unannotated residues; the reported precision metrics therefore likely represent a lower bound. Nonetheless, notable performance differences were observed across classes. For N-glycosylation, ViralMap achieved higher precision (0.648 vs. 0.270) and higher recall (0.913 vs. 0.827) than NetNGlyc. Notably, only 26% of canonical Asn-Xaa-Ser/Thr motifs in the dataset are annotated as glycosylated. When evaluated exclusively on these canonical motifs, ViralMap maintained similar performance, yielding a mean precision of 0.671 and a recall of 0.918 **(Supplementary Table 4)**. This pattern suggests that ViralMap leverages contextual sequence features to inform glycosylation predictions rather than strictly relying on motif detection.

We observed a similar dynamic with furin cleavage site prediction, where only 3.2% of canonical Arg-Xaa-Lys/Arg-Arg motifs are annotated as cleaved in the database. Across all residues, ViralMap achieved a mean precision of 0.445 and a recall of 0.780. When stratifying the evaluation by canonical motif context, performance remained consistent (mean precision 0.502; recall 0.792) **(Supplementary Table 5)**. While the high cross-fold variance in ViralMap furin performance highlights the inherent difficulty of generalizing from a highly sparse dataset, these results suggest that the model’s predictions are not solely dictated by the presence of canonical furin cleavage motifs.

This apparent contextual prediction also extended to disulfide bond sites and chain cleavage sites. Disulfide bond prediction achieved strong performance. Only 11.6% of cysteines in the dataset are annotated as participating in disulfide bonds (Supplementary Table 6), yet the model selectively identified bonding cysteines with a precision of 0.545 and recall of 0.869 when evaluated exclusively on cysteine residues **(Supplementary Table 6).** This suggests the model incorporates local sequence context to distinguish between bonding and non-bonding cysteines rather than acting as a simple cysteine presence scanner. Similarly, chain cleavage, a class also lacking a comparable sequence-based benchmark, achieved the second-highest PR-AUC (0.895). As detailed in **Supplementary Table 7,** performance was stronger on single-chain proteins (precision 0.922, recall 0.964) than on multi-chain proteins (precision 0.784, recall 0.593), consistent with the greater complexity of predicting internal processing sites relative to terminal boundaries.

For topology and localization, ViralMap performed comparably to DeepTMHMM, a highly specialized tool for topological prediction. ViralMap yielded higher precision and recall for signal peptides. For transmembrane and cytoplasmic domains, performance trade-offs were observed: compared to DeepTMHMM, ViralMap achieved higher recall but lower precision for transmembrane domains (recall: 0.868 vs. 0.695; precision: 0.884 vs. 0.941), whereas the reverse was true for cytoplasmic domains. Extracellular domain predictions remained highly comparable between the two tools.

Among structural features, intrinsically disordered regions (IDRs) achieved the highest overall PR-AUC (0.942). While AIUPred achieved near-complete recall for IDRs (0.993), ViralMap provided a more balanced predictive profile with higher precision (0.832 vs. 0.388). Finally, while coiled coils remained a quantitatively difficult class to predict globally for both ViralMap and DeepCoil, the model nevertheless demonstrated a strong capacity to localize functionally critical coiled coil motifs within canonical viral fusion machinery, a capability we explore further in the subsequent case studies.

Taken together, these results demonstrate that ViralMap achieves performance that is competitive with or exceeds dedicated single-task tools across the eight benchmarked annotation classes within a single unified model. The remaining two classes, chain cleavage sites and disulfide bond sites, for which no comparable tools requiring only primary sequence as input were available, showed promising standalone performance.

### Case studies: viral glycoprotein annotation

To demonstrate ViralMap’s predictive capabilities, we visualize the annotation profiles of two viral antigens: the SARS-CoV-2 spike glycoprotein **(Fig. 3a, 3b)** and the HIV-1 envelope (Env, gp160) glycoprotein **(Fig. 3c).** Both glycoproteins were held out during training according to the cross-validation approach described in **Section 2.3**, in which folds were assigned using a viral-family-aware strategy: large families containing more than 200 proteins were distributed across multiple folds, while smaller families were allocated to individual folds **(Supplementary Fig. 6).** This design enables two complementary tests of generalization. The *Retroviridae* family was distributed across folds, so the HIV-1 gp160 prediction provides a test of cross-strain generalization at the homology clustering thresholds defined in Section 2.3. The *Coronaviridae* family, comprising fewer than 200 proteins, was allocated to a single fold, making the spike protein prediction a more stringent test of novel-family generalization. Furthermore, because the SARS-CoV-2 spike protein (1,273 residues) exceeded the sequence-length filter applied during data curation (Section 2.1), it was entirely absent from the training data. Inference on sequences beyond this length is possible because ESM-2 employs rotary positional encodings, which generalize to lengths not seen during fine-tuning.

The SARS-CoV-2 spike protein is a trimeric fusion protein. ViralMap recovered its overarching topological pattern, identifying the N-terminal signal peptide, extracellular domain, transmembrane domain, and C-terminal cytoplasmic region **(Fig. 3b, Topology & Localization)**. To facilitate host-cell fusion, the spike protein undergoes significant proteolytic processing^51,52^ to separate its receptor-binding (N-terminal) S1 subunit from its membrane fusion (C-terminal) S2 subunit. ViralMap identified both the primary furin cleavage site (R685/S686) at the S1/S2 boundary and a secondary protease cleavage site (S2’, R815/S816) **(Fig. 3b, Post-translational Modifications)**. The S2 subunit undergoes further conformational change to drive host-cell membrane fusion, a process mediated by two coiled-coil regions: heptad repeat 1 (HR1) and heptad repeat 2 (HR2). Both HR1 and HR2 have been the central focus of recombinant subunit vaccines.^53,54^ ViralMap localized both domains **(Fig. 3a**; **Fig. 3b, Structural Features)**, suggesting the model can identify functional fusion machinery solely from primary sequence. Beyond these core structural components, ViralMap captured all 22 reference N-glycosylation sites comprising the spike protein’s glycan shield, which plays a significant role in escaping antibody recognition.^55^ The model also correctly identified the 30 cysteines participating in the extracellular disulfide bond network **(Fig. 3b, Post-translational Modifications)**, with minor over-prediction at two non-reference cysteines (C1243 and C1253). Notably, we observed that despite strong global performance for the IDR class **(Table 1)**, the model failed to resolve the spike protein’s disordered regions **(Fig. 3b, Structural Features)**, likely due to the challenge of generalizing to novel viral families.

We observed a similarly detailed annotation profile when processing HIV-1 gp160 with ViralMap. Like the spike protein, gp160 is a trimeric fusion protein consisting of two primary subunits: gp120 and gp41.^56^ ViralMap characterized the topology of gp160 **(Fig. 3c, Topology & Localization)** and identified the furin cleavage site (R511/S512) that separates the gp120 and gp41 subunits. All 29 reference N-glycosylation sites forming the dense glycan shield on gp160 were identified, and 18 out of 20 cysteines involved in disulfide bonds were identified **(Fig. 3c, Post-translational Modifications)**. The model also localized key structural features in the gp41 subunit **(Fig. 3c, Structural Features)**, including a coiled-coil domain in the N-terminal region (residues 633-667) that plays a critical role in membrane fusion.^57^ Furthermore, ViralMap identified an intrinsically disordered region annotated in UniProt (reference residues 718-742), which is consistent with predicted disorder profiles of gp41s from 50 different HIV-1 samples.^58^

## DISCUSSION

Developing effective vaccines often requires targeted sequence changes to viral proteins to elicit protective immune responses. Determining which modifications to make depends heavily on the unique biology of the target virus. This necessitates a comprehensive, accurate annotation profile to inform experimentalists’ decisions. As pandemic preparedness initiatives such as the CEPI 100 Days Mission demand increasingly rapid responses to novel pathogens, the ability to generate these profiles directly from newly sequenced viruses becomes essential. While general protein annotation tools are widely available, viral proteins present distinct profiling challenges due to their rapid evolution and distance from the eukaryotic proteins typically used to train existing models, and assembling multi-tool pipelines to obtain comprehensive annotations remains technically demanding. To address this, we developed ViralMap to meet three primary objectives: providing simultaneous multi-label prediction for features relevant to antigen design, specializing explicitly in eukaryotic viral sequences, achieving performance competitive with existing specialized annotation tools.

Our results demonstrate that ViralMap successfully accomplishes these three goals. By leveraging the pretrained representations of the ESM-2 language model with a custom post-processing pipeline, ViralMap effectively classifies ten distinct annotation classes spanning topology and localization, post-translational modifications, and structural features. As evidenced by the overall classification performance **(Table 1)** and individual antigen-profiling case studies **(Fig. 3)**, the model can directly extract these diverse annotations from primary sequences, providing a unified alternative to disjointed multi-tool pipelines. Importantly, our benchmarking comparison was conducted specifically on eukaryotic viral proteins. The established tools evaluated alongside ViralMap were developed for broader protein annotation tasks, and their performance on general or non-viral datasets may differ from the results reported here. Our comparison therefore reflects the specific use case that ViralMap was designed to address rather than a general assessment of these tools’ capabilities.

The design and evaluation of ViralMap necessitated pragmatic trade-offs. Given the severe sparsity and inconsistent quality of viral annotations, our dataset curation pipeline prioritized entries with high-quality individual annotations while retaining instances of rare annotation types. Because ViralMap predicts ten classes simultaneously, this curation could not rely solely on sequence-based redundancy reduction and instead required an annotation-aware selection strategy that accounted for quality and coverage across classes. Selecting UniProt entries in an annotation-first manner, however, naturally introduced taxonomic biases **(Supplementary Fig. 6)**. Future iterations of the dataset curation process could improve upon this by stratifying selection within viral families to ensure a more balanced taxonomic distribution. Furthermore, addressing false-negative annotations and mitigating the high-performance variance across cross-validation folds for certain classes will require expanding our training sets. This could involve aggregating curated data across multiple specialized databases or introducing synthetic examples. Future versions of the model will also aim to broaden annotation capabilities, such as predicting explicit disulfide bond pairings.

Architectural constraints define additional avenues for future development. Currently, ViralMap enforces biological rules locally by using separate HMMs to manage mutually exclusive states within specific categories, such as topology. While this design preserves the base model’s strong predictive capabilities, subsequent iterations could transition toward a unified decoding framework. By defining a joint state space across all target classes, future models could enforce a more global biological grammar. Ultimately, ViralMap bridges a critical gap in the rapid-response pipeline for emerging viral threats. By converting primary protein sequences into comprehensive annotation profiles in a single pass, it provides a practical and scalable link between the extraction of viral sequences from newly characterized genomes and the functional information required for downstream antigen engineering.

## CODE AVAILABILITY

ViralMap is implemented in Python and will be available as a command-line interface on GitHub at https://github.com/HMRI-ADAPT/vmap. A web interface will be accessible at the Disease X Knowledge Base (dxkb.org).

## ACKNOWLEDGMENTS

We thank ADAPT members Dan Boutz, Thomas Segall-Shapiro, and Raghav Schroff for helpful discussions over the course of the study. This work was funded in part by the CEPI Immunogen Design for Disease X program (JDG) and the ARPA-H APECx SHIELD consortium (JDG). The research was supported in part by the Houston Methodist Academic Institute Infectious Diseases Fund and many generous Houston philanthropists (JDG). We are grateful to Carole Walter Looke, Jim Looke, and Evan H Katz for their generous philanthropic gift to ADAPT (JDG). The funders had no role in the design and conduct of the study; collection, management, analysis, and interpretation of the data; preparation, review, or approval of the manuscript; and decision to submit the manuscript for publication.

## AUTHOR CONTRIBUTIONS

Conceptualization: SD, APH, JDG; Investigation & visualization: SD, JDG; Methodology: all authors; Software: SD; Data Curation: SD; Writing – Original draft: SD; Writing – Reviewing & Editing: all authors; Supervision: APH, JDG

## REFERENCES

1. Pollard, A. J. & Bijker, E. M. A guide to vaccinology: from basic principles to new developments. Nat. Rev. Immunol. 21, 83–100 (2021).

2. Essalmani, R. et al. Distinctive Roles of Furin and TMPRSS2 in SARS-CoV-2 Infectivity. J. Virol. 96, e00128–22 (2022).

3. Byrne, P. O. & McLellan, J. S. Principles and practical applications of structure-based vaccine design. Curr. Opin. Immunol. 77, 102209 (2022).

4. Hsieh, C.-L. et al. Structure-based design of prefusion-stabilized SARS-CoV-2 spikes. Science 369, 1501–1505 (2020).

5. McLellan, J. S. et al. Structure-Based Design of a Fusion Glycoprotein Vaccine for Respiratory Syncytial Virus. Science 342, 592–598 (2013).

6. Crooks, E. T. et al. Vaccine-Elicited Tier 2 HIV-1 Neutralizing Antibodies Bind to Quaternary Epitopes Involving Glycan-Deficient Patches Proximal to the CD4 Binding Site. PLOS Pathog. 11, e1004932 (2015).

7. Zhou, T. et al. Quantification of the Impact of the HIV-1-Glycan Shield on Antibody Elicitation. Cell Rep. 19, 719–732 (2017).

8. Imai, K. & Nakai, K. Tools for the Recognition of Sorting Signals and the Prediction of Subcellular Localization of Proteins From Their Amino Acid Sequences. Front. Genet. 11, 607812 (2020).

9. Upadhyay, C. et al. Signal peptide exchange alters HIV-1 envelope antigenicity and immunogenicity. Front. Immunol. 15, 1476924 (2024).

10. Barrett, A. J., Rawlings, N. D. & Woessner, J. F. (eds.) Handbook of Proteolytic Enzymes 3rd edn (Academic Press, 2013).

11. Wu, Y. & Zhao, S. Furin cleavage sites naturally occur in coronaviruses. Stem Cell Res. 50, 102115 (2021).

12. Esmail, S. & Manolson, M. F. Advances in understanding N-glycosylation structure, function, and regulation in health and disease. Eur. J. Cell Biol. 100, 151186 (2021).

13. Vigerust, D. J. & Shepherd, V. L. Virus glycosylation: role in virulence and immune interactions. Trends Microbiol. 15, 211–218 (2007).

14. Matthews, J. M., Young, T. F., Tucker, S. P. & Mackay, J. P. The Core of the Respiratory Syncytial Virus Fusion Protein Is a Trimeric Coiled Coil. J. Virol. 74, 5911–5920 (2000).

15. Watanabe, S. et al. Functional Importance of the Coiled-Coil of the Ebola Virus Glycoprotein. J. Virol. 74, 10194–10201 (2000).

16. Mughal, F. & Caetano-Anollés, G. Evolution of intrinsic disorder in the structural domains of viral and cellular proteomes. Sci. Rep. 15, 2878 (2025).

17. Baker, R. E. et al. Infectious disease in an era of global change. Nat. Rev. Microbiol. 20, 193–205 (2022).

18. Coalition for Epidemic Preparedness Innovations (CEPI). The 100 Days Mission. https://cepi.net/100-days-mission (accessed 3 April 2026).

19. Barnsley, G. et al. Impact of the 100 days mission for vaccines on COVID-19: a mathematical modelling study. Lancet Glob. Health 12, e1764–e1774 (2024).

20. Prakash, H. et al. Wastewater sequencing reveals persistent circulation and rising prevalence of several oncogenic viruses across Texas. Preprint at 10.1101/2025.09.17.25335998 (2025).

21. Tisza, M. J., Petrosino, J. F. & Javornik Cregeen, S. J. Cenote-Taker 3 for fast and accurate virus discovery and annotation of the virome. Preprint at 10.1101/2025.08.20.671380 (2025).

22. Dumitrescu, A. et al. TSignal: a transformer model for signal peptide prediction. Bioinformatics 39, i347–i356 (2023).

23. Teufel, F. et al. SignalP 6.0 predicts all five types of signal peptides using protein language models. Nat. Biotechnol. 40, 1023–1025 (2022).

24. Lotthammer, J. M., Ginell, G. M., Griffith, D., Emenecker, R. J. & Holehouse, A. S. Direct prediction of intrinsically disordered protein conformational properties from sequence. Nat. Methods 21, 465–476 (2024).

25. Redl, I., et al. ADOPT: intrinsic protein disorder prediction through deep bidirectional transformers. NAR Genomics Bioinforma. 5, lqad041 (2023).

26. Hou, X., Wang, Y., Bu, D., Wang, Y. & Sun, S. EMNGly: predicting N-linked glycosylation sites using the language models for feature extraction. Bioinformatics 39, btad650 (2023).

27. Pitti, T. et al. N-GlyDE: a two-stage N-linked glycosylation site prediction incorporating gapped dipeptides and pattern-based encoding. Sci. Rep. 9, 15975 (2019).

28. Bernhofer, M. & Rost, B. TMbed: transmembrane proteins predicted through language model embeddings. BMC Bioinformatics 23, 326 (2022).

29. Hallgren, J. et al. DeepTMHMM predicts alpha and beta transmembrane proteins using deep neural networks. Preprint at 10.1101/2022.04.08.487609 (2022).

30. Nomburg, J. et al. Birth of protein folds and functions in the virome. Nature 633, 710–717 (2024).

31. Sawhney, R. et al. Fine-tuning protein language models unlocks the potential of underrepresented viral proteomes. PeerJ 13, e19919 (2025).

32. Pearson, W. R. An Introduction to Sequence Similarity (“Homology”) Searching. Curr. Protoc. Bioinforma. 42, (2013).

33. Fang, Z., Feng, T., Zhou, H. & Chen, M. DeePVP: Identification and classification of phage virion proteins using deep learning. GigaScience 11, giac076 (2022).

34. Flamholz, Z. N., Biller, S. J. & Kelly, L. Large language models improve annotation of prokaryotic viral proteins. Nat. Microbiol. 9, 537–549 (2024).

35. Shang, J., Peng, C., Tang, X. & Sun, Y. PhaVIP: Phage VIrion Protein classification based on chaos game representation and Vision Transformer. Bioinformatics 39, i30–i39 (2023).

36. Flamholz, Z. N., Li, C. & Kelly, L. Improving viral annotation with artificial intelligence. mBio 15, e03206–23 (2024).

37. Freeman, K. G. et al. Virion glycosylation influences mycobacteriophage immune recognition. Cell Host Microbe 31, 1216–1231.e6 (2023).

38. Cui, Y., Jia, M., Lin, T.-Y., Song, Y. & Belongie, S. Class-Balanced Loss Based on Effective Number of Samples. Preprint at 10.48550/arXiv.1901.05555 (2019).

39. Loshchilov, I. & Hutter, F. Decoupled Weight Decay Regularization. Preprint at 10.48550/arXiv.1711.05101 (2019).

40. Ludwiczak, J., Winski, A., Szczepaniak, K., Alva, V. & Dunin-Horkawicz, S. DeepCoil—a fast and accurate prediction of coiled-coil domains in protein sequences. Bioinformatics 35, 2790–2795 (2019).

41. Erdős, G. & Dosztányi, Z. AIUPred: combining energy estimation with deep learning for the enhanced prediction of protein disorder. Nucleic Acids Res. 52, W176–W181 (2024).

42. Gupta, R. & Brunak, S. Prediction of glycosylation across the human proteome and the correlation to protein function. in Biocomputing 2002 310–322 (WORLD SCIENTIFIC, Kauai, Hawaii, USA, 2001). doi:10.1142/9789812799623_0029.

43. Duckert, P., Brunak, S. & Blom, N. Prediction of proprotein convertase cleavage sites. Protein Eng. Des. Sel. 17, 107–112 (2004).

44. Savojardo, C. et al. Improving the prediction of disulfide bonds in Eukaryotes with machine learning methods and protein subcellular localization. Bioinformatics 27, 2224–2230 (2011).

45. Lin, Z. et al. Evolutionary-scale prediction of atomic-level protein structure with a language model.

46. Ito, J. et al. A Protein Language Model for Exploring Viral Fitness Landscapes. Preprint at 10.1101/2024.03.15.584819 (2024).

47. Hie, B., Zhong, E. D., Berger, B. & Bryson, B. Learning the language of viral evolution and escape.

48. The UniProt Consortium et al. UniProt: the Universal Protein Knowledgebase in 2025. Nucleic Acids Res. 53, D609–D617 (2025).

49. Steinegger, M. & Söding, J. MMseqs2 enables sensitive protein sequence searching for the analysis of massive data sets. Nat. Biotechnol. 35, 1026–1028 (2017).

50. Nadendla, S. et al. ECO: the Evidence and Conclusion Ontology, an update for 2022. Nucleic Acids Res. 50, D1515–D1521 (2022).

51. Cai, Y. et al. Distinct conformational states of SARS-CoV-2 spike protein.

52. Wrapp, D. et al. Cryo-EM structure of the 2019-nCoV spike in the prefusion conformation. Science 367, 1260–1263 (2020).

53. Pang, W. et al. A variant-proof SARS-CoV-2 vaccine targeting HR1 domain in S2 subunit of spike protein. Cell Res. 32, 1068–1085 (2022).

54. Lu, Y., et al. Recombinant protein HR212 targeting heptad repeat 2 domain in spike protein S2 subunit elicits broad-spectrum neutralizing antibodies against SARS-CoV-2 and its variants. MedComm 6, e70088 (2025).

55. Grant, O. C., Montgomery, D., Ito, K. & Woods, R. J. Analysis of the SARS-CoV-2 spike protein glycan shield reveals implications for immune recognition. Sci. Rep. 10, 14991 (2020).

56. Arrildt, K. T., Joseph, S. B. & Swanstrom, R. The HIV-1 Env Protein: A Coat of Many Colors. Curr. HIV/AIDS Rep. 9, 52–63 (2012).

57. Mo, H. et al. Conserved residues in the coiled-coil pocket of human immunodeficiency virus type 1 gp41 are essential for viral replication and interhelical interaction. Virology 329, 319–327 (2004).

58. Xue, B., Mizianty, M. J., Kurgan, L. & Uversky, V. N. Protein intrinsic disorder as a flexible armor and a weapon of HIV-1. Cell. Mol. Life Sci. 69, 1211–1259 (2012).

