## Supplementary Information for "ViralMap: Predicting Features in Viral Proteins from Primary Sequence"

### Supplementary Figures

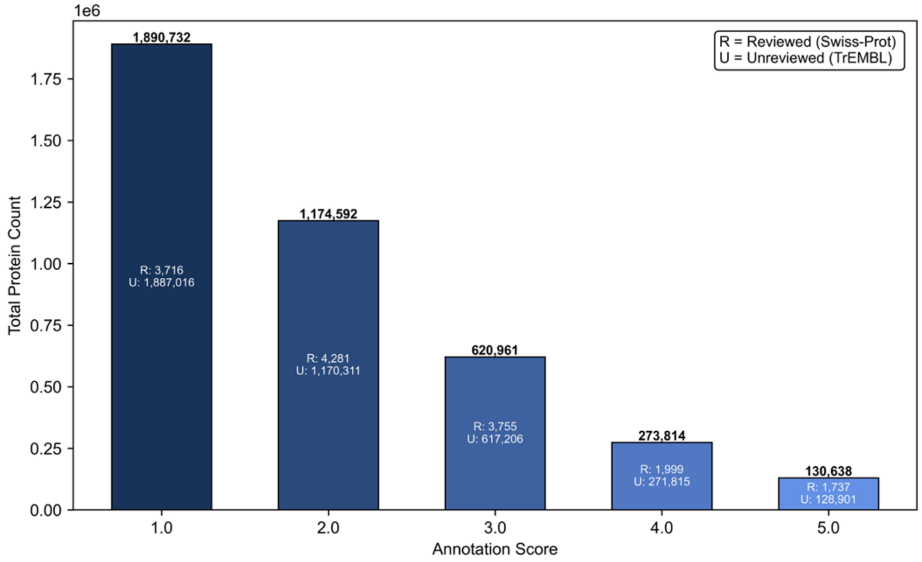


**Supplementary Figure 1**. The initial download of non-phage viral proteins (n=4,090,737 proteins) from UniProt was composed mainly of poorly annotated entries, with most sequences assigned UniProt annotation scores of 3.0 or lower. The Swiss-Prot (reviewed) designation did not necessarily indicate that the protein was well annotated, since most proteins with a 5.0 annotation score in the dataset were TrEMBL (unreviewed) records.

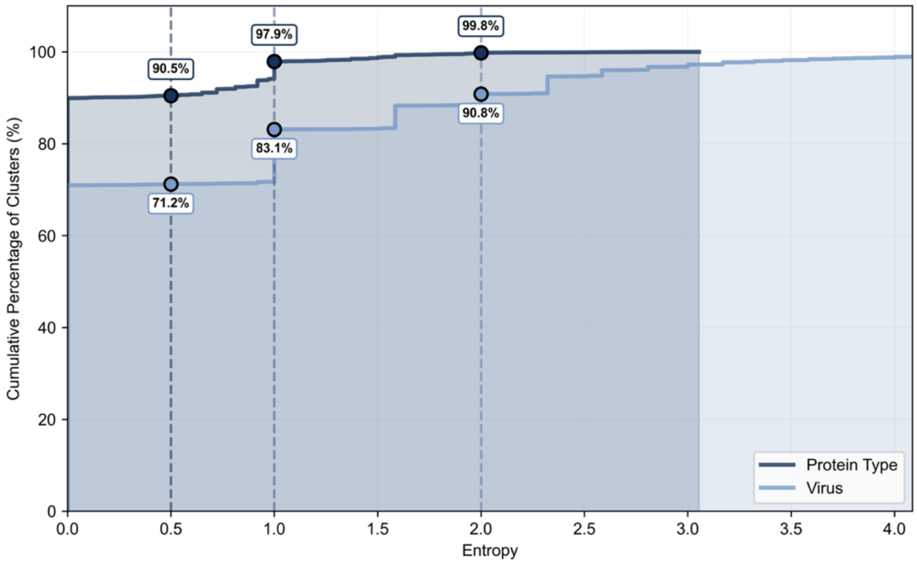


**Supplementary Figure 2.** MMSeqs2 clustering (60% sequence identity and 50% bidirectional coverage) produces largely homogeneous protein-type (e.g., glycoprotein, matrix protein, etc.) and virus-level downselection clusters. Shown are cumulative distributions of Shannon entropy for UniProt protein names and UniProt organism (virus). The entropy metric shows how similar UniProt entries are within a given cluster with respect to protein type and virus, on a scale from 0 to infinity. For protein type, 99.8% of all clusters have entropy ≤ 2.0, indicating strong homogeneity in protein types. Virus-level homogeneity was similarly high, with 90.8% of clusters having entropy ≤ 2.0. These results suggest that most clusters group variants of the same viral protein, enabling representative selection as described in **Supplementary Note 1**.

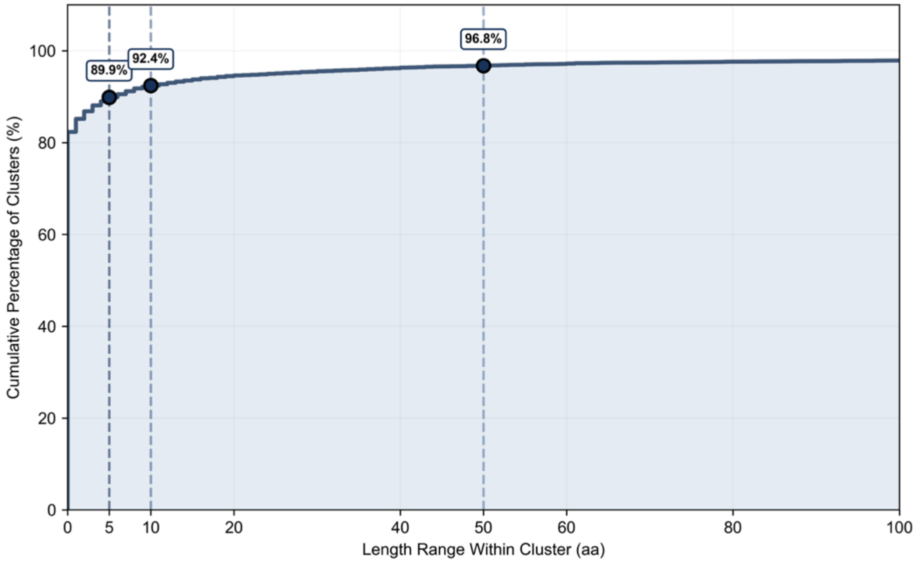


**Supplementary Figure 3.** Intra-cluster protein length variation is minimal for the majority of MMSeqs2 downselection clusters. Shown is the cumulative distribution of sequence length range (max length – min length) within clusters. Most clusters exhibited narrow length variation, with 89.9% having a length range ≤5 amino acids and 96.8% ≤ 50 amino acids. This indicates that clustered proteins are mostly consistent in length, supporting their use as interchangeable candidates during representative selection **(Supplementary Note 1)**.

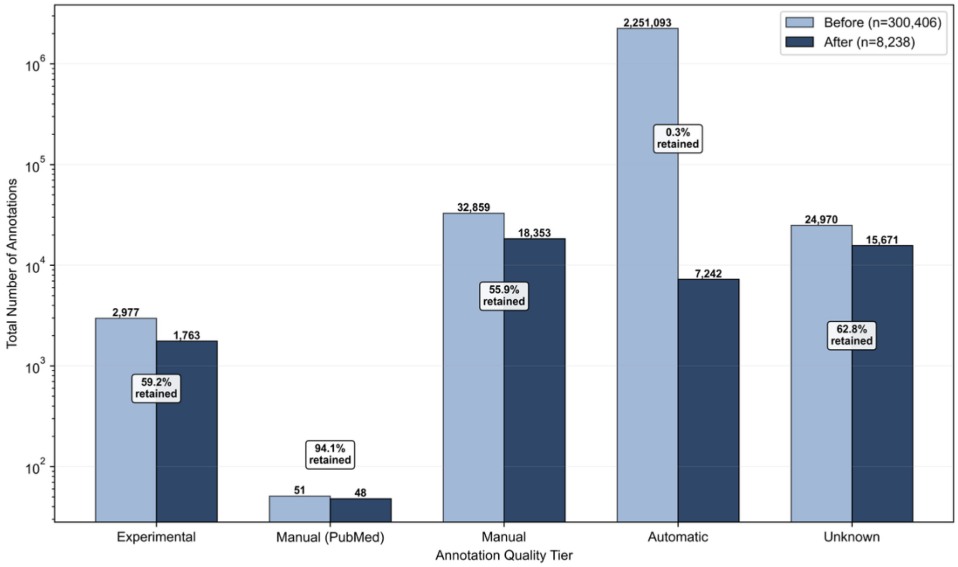


**Supplementary Figure 4.** Annotation retention by evidence tier following representative selection via Algorithm 1 **(Supplementary Note 1)**. Grouped bars show annotation counts before (n=300,406 proteins) and after (n=8,238 proteins) selection. Retention rates varied by evidence quality: experimental (59.2%), manual with PubMed (94.1%), manual (55.9%), automatic (0.3%), and unknown (62.8%). The algorithm successfully prioritized selecting proteins with higher-quality annotations across all classes while reducing redundancy and preferentially eliminating proteins that contributed automatic annotations.

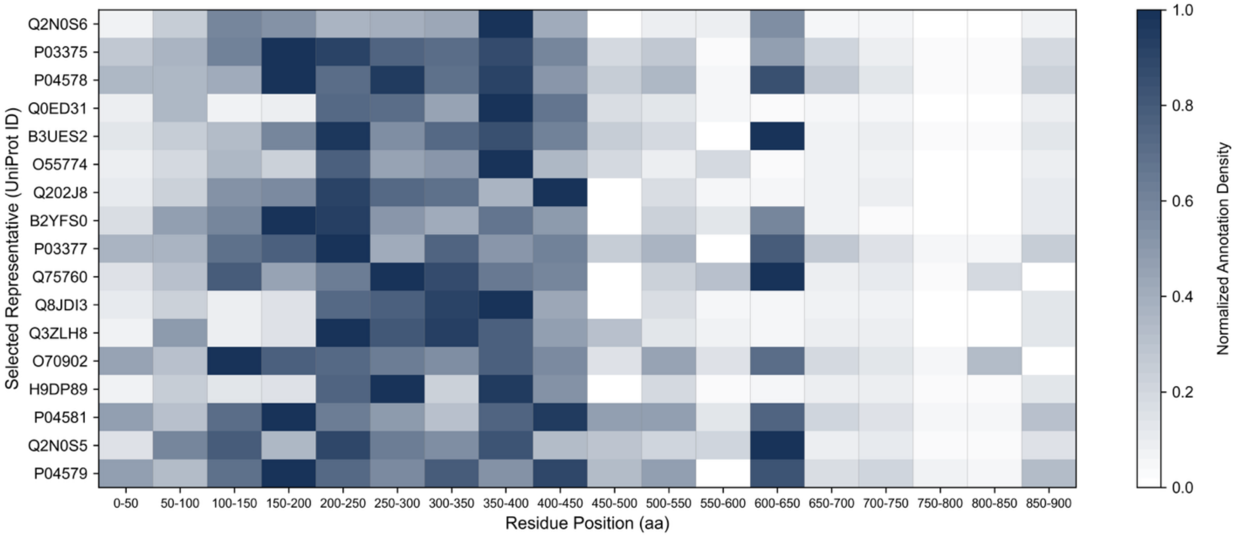


**Supplementary Figure 5.** Annotation coverage across selected HIV-1 envelope protein representatives. The heatmap shows combined annotation density across all 10 classes for sequence bins (50 residues each) of the 17 representatives retained from 60,348 original entries. Despite a substantial reduction in cluster size, selected representatives maintain both annotation density and coverage.

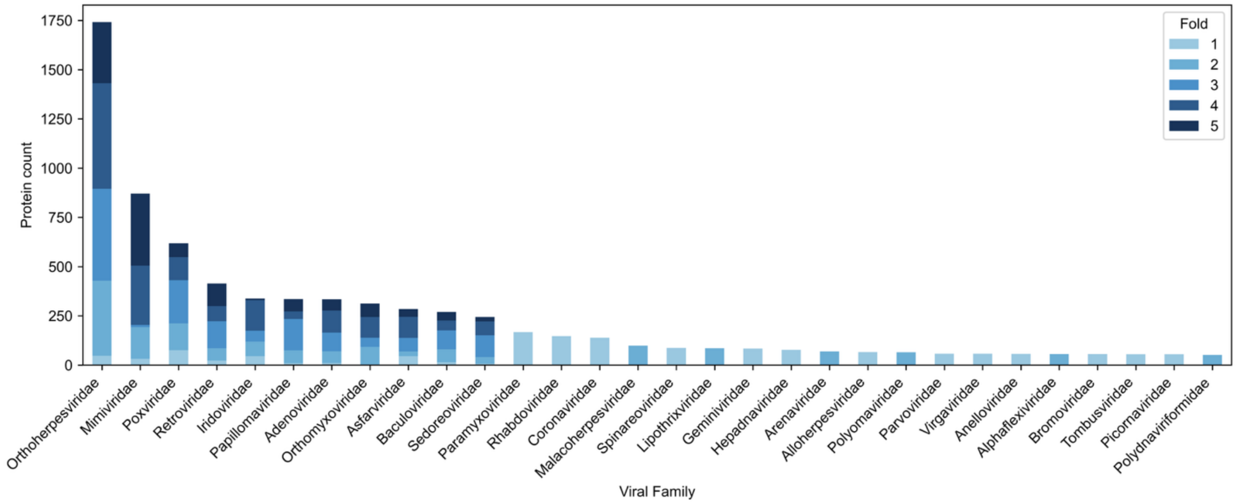


**Supplementary Figure 6.** Viral family distribution across fold assignments in the final ViralMap dataset highlights the top 30 families by protein count, which together account for over 88% of the 97 unique families in the dataset. Fold assignments aimed to balance annotation class distributions at the residue level and enhance taxonomic diversity. Large families (with protein counts over 200) were divided into separate folds to minimize taxonomic bias, while smaller families were grouped into different folds to increase diversity in model evaluation. To prevent homology leakage, all clusters were kept intact regardless of family size (no clusters were split across folds).

### Supplementary Tables

| ECO Code | UniProt Description (Abbreviated) | Sources | Type | Score |
|---|---|---|---|---|
| ECO:0000269 | Manually curated information with published experimental evidence | PubMed, Ref.* | Experimental | 4 |
| ECO:0007744 | Combinatorial evidence from experimental and computational sources; manual assertion | PDB | Experimental | 4 |
| ECO:0007829 | Combinatorial evidence from experimental and computational sources; automatic assertion | PDB | Experimental | 4 |
| ECO:0000305 | Curator inference based on scientific knowledge or article content | PubMed, None | Manual (PubMed) | 3 |
| ECO:0000250 | Propagated from related experimentally characterized protein | None, UniProtKB | Manual | 2 |
| ECO:0000255 | Sequence model match evidence; curator-verified computational prediction | HAMAP-Rule, None, PROSITE-ProRule | Manual | 2 |
| ECO:0000256 | Automatic assertion generated by UniProtKB annotation system | HAMAP-Rule, SAM, RuleBase, PIRSR | Automatic | 1 |
| - | - | - | Unknown | 0.5 |

**Supplementary Table 1.** Mapping of Evidence and Conclusion Ontology (ECO) codes in UniProt annotations to annotation scores to select cluster representatives and assess dataset quality **(Supplementary Note 1, Supplementary Table 2)**. After the initial filtering, which resulted in 300,406 proteins (Section 2.1), 7 unique ECO codes were observed across all annotations in the dataset. By combining the ECO code of a given annotation with the underlying source, we categorized all annotations as belonging to 5 categories: Experimental, Manual (PubMed), Manual, Automatic, or Unknown. All annotations with ECO codes ECO:0007744 and ECO:0007829 originated from experimental PDB entries, which we confirmed by referencing each to the RCSB PDB. ECO codes ECO:0000269, ECO:0007744, and ECO:0007829 were therefore classified as experimental annotations. ECO code ECO:0000305 were categorized as Manual or Manual (PubMed), depending on whether UniProt listed a publication reference with the annotation.

| Class | Fold | Experimental | Manual (PubMed) | Manual | Automatic | Unknown | Total |
|---|---|---|---|---|---|---|---|
| SP | Fold 1 | 4 / 87 | 1 / 19 | 169 / 3,867 | – | 6 / 120 | 180 / 4,093 |
| | Fold 2 | 5 / 126 | 1 / 62 | 145 / 3,774 | – | 1 / 21 | 152 / 3,983 |
| | Fold 3 | 8 / 170 | – | 160 / 3,871 | – | 2 / 20 | 170 / 4,061 |
| | Fold 4 | 1 / 31 | – | 165 / 3,921 | – | 4 / 105 | 170 / 4,057 |
| | Fold 5 | 3 / 142 | – | 166 / 3,849 | – | 6 / 74 | 175 / 4,065 |
| | **Total** | **21 / 556** | **2 / 81** | **805 / 19,282** | – | **19 / 340** | **847 / 20,259** |
| CC | Fold 1 | 1 / 66 | – | 135 / 4,842 | 3 / 102 | – | 139 / 5,010 |
| | Fold 2 | – | – | 118 / 4,433 | 10 / 489 | – | 128 / 4,922 |
| | Fold 3 | 8 / 366 | – | 99 / 4,096 | 7 / 240 | 1 / 46 | 115 / 4,748 |
| | Fold 4 | – | – | 83 / 3,717 | 38 / 1,225 | – | 121 / 4,942 |
| | Fold 5 | – | – | 112 / 4,551 | 13 / 447 | – | 125 / 4,998 |
| | **Total** | **9 / 432** | – | **547 / 21,639** | **71 / 2,503** | **1 / 46** | **628 / 24,620** |
| TM | Fold 1 | 13 / 294 | 2 / 42 | 603 / 12,768 | 33 / 722 | – | 651 / 13,826 |
| | Fold 2 | 1 / 16 | – | 578 / 12,084 | 34 / 827 | – | 613 / 12,927 |
| | Fold 3 | 1 / 26 | – | 552 / 11,691 | 93 / 2,034 | 1 / 29 | 647 / 13,780 |
| | Fold 4 | – | – | 577 / 12,253 | 83 / 2,010 | – | 660 / 14,263 |
| | Fold 5 | – | 1 / 18 | 600 / 12,678 | 40 / 994 | 1 / 24 | 642 / 13,714 |
| | **Total** | **15 / 336** | **3 / 60** | **2,910 / 61,474** | **283 / 6,587** | **2 / 53** | **3,213 / 68,510** |
| NG | Fold 1 | 75 / 75 | 4 / 4 | 1,149 / 1,149 | 2 / 2 | 5 / 5 | 1,235 / 1,235 |
| | Fold 2 | 66 / 66 | 2 / 2 | 1,308 / 1,308 | 153 / 153 | – | 1,529 / 1,529 |
| | Fold 3 | 92 / 92 | – | 1,134 / 1,134 | 8 / 8 | – | 1,234 / 1,234 |
| | Fold 4 | 296 / 296 | – | 1,025 / 1,025 | – | – | 1,321 / 1,321 |
| | Fold 5 | 45 / 45 | 2 / 2 | 1,359 / 1,359 | – | 1 / 1 | 1,407 / 1,407 |
| | **Total** | **574 / 574** | **8 / 8** | **5,975 / 5,975** | **163 / 163** | **6 / 6** | **6,726 / 6,726** |
| DR | Fold 1 | 1 / 23 | 3 / 89 | 4 / 92 | 1,125 / 50,733 | – | 1,133 / 50,937 |
| | Fold 2 | 2 / 33 | 3 / 85 | – | 1,032 / 51,468 | – | 1,037 / 51,586 |
| | Fold 3 | 1 / 23 | – | 8 / 490 | 874 / 49,770 | – | 883 / 50,283 |
| | Fold 4 | – | – | – | 967 / 50,561 | – | 967 / 50,561 |
| | Fold 5 | – | – | – | 966 / 46,724 | – | 966 / 46,724 |
| | **Total** | **4 / 79** | **6 / 174** | **12 / 582** | **4,964 / 249,256** | – | **4,986 / 250,091** |
| FR | Fold 1 | – | – | 20 / 40 | – | 1 / 2 | 21 / 42 |
| | Fold 2 | 1 / 2 | – | 18 / 36 | – | 1 / 2 | 20 / 40 |
| | Fold 3 | 2 / 4 | – | 18 / 36 | – | – | 20 / 40 |
| | Fold 4 | – | – | 9 / 18 | 11 / 22 | – | 20 / 40 |
| | Fold 5 | – | – | 14 / 28 | 6 / 12 | – | 20 / 40 |
| | **Total** | **3 / 6** | – | **79 / 158** | **17 / 34** | **2 / 4** | **101 / 202** |
| CH | Fold 1 | – | – | 391 / 391 | 12 / 12 | 3,445 / 3,445 | 3,848 / 3,848 |
| | Fold 2 | – | 4 / 4 | 587 / 587 | 266 / 266 | 2,982 / 2,982 | 3,839 / 3,839 |
| | Fold 3 | 2 / 2 | – | 611 / 611 | 129 / 129 | 2,919 / 2,919 | 3,661 / 3,661 |
| | Fold 4 | 2 / 2 | – | 419 / 419 | 496 / 496 | 2,762 / 2,762 | 3,679 / 3,679 |
| | Fold 5 | – | – | 397 / 397 | 57 / 57 | 2,694 / 2,694 | 3,148 / 3,148 |
| | **Total** | **4 / 4** | **4 / 4** | **2,405 / 2,405** | **960 / 960** | **14,802 / 14,802** | **18,175 / 18,175** |
| DB | Fold 1 | 214 / 214 | 1 / 1 | 1,258 / 1,258 | 27 / 27 | 2 / 2 | 1,502 / 1,502 |
| | Fold 2 | 393 / 393 | 3 / 3 | 1,007 / 1,007 | 110 / 110 | 21 / 21 | 1,534 / 1,534 |
| | Fold 3 | 292 / 292 | 4 / 4 | 1,141 / 1,141 | 14 / 14 | 13 / 13 | 1,464 / 1,464 |
| | Fold 4 | 617 / 617 | – | 683 / 683 | 285 / 285 | 6 / 6 | 1,591 / 1,591 |
| | Fold 5 | 513 / 513 | 2 / 2 | 944 / 944 | 115 / 115 | 19 / 19 | 1,593 / 1,593 |
| | **Total** | **2,029 / 2,029** | **10 / 10** | **5,033 / 5,033** | **551 / 551** | **61 / 61** | **7,684 / 7,684** |
| CY | Fold 1 | 8 / 467 | 3 / 181 | 255 / 15,478 | 5 / 155 | – | 271 / 16,281 |
| | Fold 2 | 3 / 116 | 5 / 303 | 295 / 12,509 | 36 / 1,166 | 4 / 32 | 343 / 14,126 |
| | Fold 3 | 1 / 24 | 1 / 42 | 286 / 11,583 | 8 / 953 | 16 / 1,512 | 312 / 14,114 |
| | Fold 4 | 3 / 352 | – | 178 / 10,715 | 11 / 1,717 | 1 / 12 | 193 / 12,796 |
| | Fold 5 | 1 / 77 | – | 257 / 12,724 | 6 / 948 | 3 / 591 | 267 / 14,340 |
| | **Total** | **16 / 1,036** | **9 / 526** | **1,271 / 63,009** | **66 / 4,939** | **24 / 2,147** | **1,386 / 71,657** |
| EX | Fold 1 | 7 / 189 | 2 / 79 | 268 / 73,011 | 2 / 413 | – | 279 / 73,692 |
| | Fold 2 | 1 / 16 | 5 / 129 | 270 / 60,552 | 19 / 662 | 4 / 576 | 299 / 61,935 |
| | Fold 3 | 1 / 499 | 1 / 235 | 301 / 58,365 | 8 / 224 | 14 / 2,050 | 325 / 61,373 |
| | Fold 4 | 4 / 249 | – | 173 / 45,372 | 1 / 652 | 2 / 492 | 180 / 46,765 |
| | Fold 5 | 1 / 10 | – | 250 / 62,377 | – | 3 / 11 | 254 / 62,398 |
| | **Total** | **14 / 963** | **8 / 443** | **1,262 / 299,677** | **30 / 1,951** | **23 / 3,129** | **1,337 / 306,163** |
| **Total** | Fold 1 | 323 / 1,415 | 16 / 415 | 4,252 / 112,896 | 1,209 / 52,166 | 3,459 / 3,574 | 9,259 / 170,466 |
| | Fold 2 | 472 / 768 | 23 / 588 | 4,326 / 96,290 | 1,660 / 55,141 | 3,013 / 3,634 | 9,494 / 156,421 |
| | Fold 3 | 408 / 1,498 | 6 / 281 | 4,310 / 93,018 | 1,141 / 53,372 | 2,966 / 6,589 | 8,831 / 154,758 |
| | Fold 4 | 923 / 1,547 | – | 3,312 / 78,123 | 1,892 / 56,968 | 2,775 / 3,377 | 8,902 / 140,015 |
| | Fold 5 | 563 / 787 | 5 / 22 | 4,099 / 98,907 | 1,203 / 49,297 | 2,727 / 3,414 | 8,597 / 152,427 |
| | **Total** | **2,689 / 6,015** | **50 / 1,306** | **20,299 / 479,234** | **7,105 / 266,944** | **14,940 / 20,588** | **45,083 / 774,087** |

**Supplementary Table 2.** Instance and residue counts stratified by evidence source and cross-validation fold for all 10 classes in the final ViralMap dataset: signal peptides (SP), coiled coils (CC), transmembrane domains (TM), N-glycosylation (NG), intrinsically disordered regions (DR), furin cleavage sites (FR), chain cleavage sites (CH), disulfide bond sites (DB), cytoplasmic domains (CY), and extracellular domains (EX). Each entry is formatted as instance count/residue count, where instance count is the number of individual examples of the annotation. For example, the dataset includes 21 signal peptides with experimental evidence.

| Tool | Classes | Raw Output | Binarization |
|---|---|---|---|
| DeepTMHMM | SP, TM, CY, EX | Per-residue topology labels: Inside (I), Membrane (M), Outside (O), Signal (S) | Direct mapping: I→CY, M→TM, O→EX, S→SP |
| DeepCoil | CC | Sharpened per-residue probabilities from peak-finding algorithm; non-peak residues assigned 0, peak residues share maximum | Contiguous regions called positive where peak maximum ≥0.5 |
| AIUPred | DR | Per-residue disorder probabilities | Residues with probability >0.5 called positive; gaps of ≤2 residues between positive regions filled |
| NetNGlyc | NG | Per-site potential scores for Asn residues in N-glycosylation motifs | Sites with potential >0.5 called positive |
| ProP | FR | Binary site calls for furin cleavage motifs | Direct (sites labeled "ProP" called positive) |

**Supplementary Table 3.** Converting benchmark tool raw outputs into binary predictions for comparison with ViralMap for eight classes: signal peptides (SP), transmembrane domains (TM), cytoplasmic domains (CY), extracellular domains (EX), coiled coils (CC), intrinsically disordered regions (DR), N-glycosylation (NG), and furin cleavage sites (FR).

| | Dataset | | | Performance (residue-level) | | | |
|---|---|---|---|---|---|---|---|
| | | | | ViralMap | | Benchmark | |
| **Context** | Total Count | N-Glycosylated? | % of Total | Precision | Recall | Precision | Recall |
| All Asparagines (Asn) | 164,751 | 6,649 | 4.04% | **0.651**±0.06 | **0.915**±0.03 | 0.270±0.02 | 0.829±0.01 |
| N-X-S/T Motifs | 25,432 | 6,622 | 26.04% | **0.671**±0.06 | **0.918**±0.03 | 0.283±0.02 | 0.832±0.01 |
| Non-Motif Asn | 139,319 | 27 | 0.02% | **0.006**±0.01 | **0.050**±0.10 | – | – |

**Supplementary Table 4**. N-glycosylation prediction performance stratified by asparagine context. Dataset columns report counts summed across 5 test folds. Precision and recall are mean ± standard deviation. Bold indicates better performance. NetNGlyc predicts on sequons by default; non-motif performance was not evaluated. Total N-glycosylation site counts are slightly lower than those in Supplementary Table 2 because counts are aggregated across cleaned test folds, where a small subset of proteins was removed **(Section 2.3)** to enforce the clustering homology threshold. Bold values indicate the better-performing method for that metric.

| | Dataset | | | Performance (residue-level) | | | |
|---|---|---|---|---|---|---|---|
| | | | | ViralMap | | Benchmark | |
| **Context** | Motif Count | Cleaved? | % of Total | Precision | Recall | Precision | Recall |
| R-X-K/R-R↓ | 2,907 | 93 | 3.2% | **0.502**±0.11 | **0.792**±0.36 | 0.051±0.01 | 0.456±0.04 |

**Supplementary Table 5.** Furin cleavage site characterization and prediction performance at the canonical motif sequence Arg-Xaa-Lys/Arg-Arg ↓, where Xaa is any amino acid, K is lysine, and the ↓ is where cleavage occurs. Dataset columns report the overall motif count summed over 5 test folds, and the count/percentage of those motifs that are annotated as cleaved (positive furin cleavage sites) in UniProt. Of the 101 total annotated furin cleavage sites in the dataset **(Supplementary Table 2)**, 93 (92%) fall within this canonical motif. Out of the remaining 8, 1 was removed from the test folds to enforce homology separation, and the remaining 7 correspond to other motifs and are excluded from the table **(Section 2.3)**. Residue-level precision and recall are computed from the two residues flanking supposed cleavage sites and are reported as mean ± standard deviation across 5 test folds. Bold values indicate the better-performing method for that metric.

| | Dataset | | | Performance (residue-level) | |
|---|---|---|---|---|---|
| **Context** | Cys Count | Disulfide? | % of Total | Precision | Recall |
| Cysteine | 64,852 | 7,522 | 11.6% | 0.545±0.07 | 0.869±0.10 |

**Supplementary Table 6.** Disulfide bond prediction performance on cysteine (Cys) residues. Dataset counts reflect the total counts across all 5 test folds. Residue-level precision and recall are reported for ViralMap disulfide bond site prediction on cysteines only as mean ± standard deviation across folds.

| | Dataset | | Performance (residue-level) | |
|---|---|---|---|---|
| | | | ViralMap | |
| **Context** | Proteins | Cleavage Sites | Precision | Recall |
| All chain proteins | 7,845 | 17,969 | 0.896±0.04 | 0.884±0.02 |
| Single chain proteins | 7,003 | 14,006 | 0.922±0.02 | 0.964±0.01 |
| Multi-chain proteins | 842 | 3,963 | 0.784±0.10 | 0.593±0.09 |

**Supplementary Table 7.** Chain cleavage site characterization and prediction performance, stratified by single-chain proteins and multi-chain proteins. Single-chain proteins have one pair of boundary positions (start and end of the mature polypeptide), while multi-chain proteins contain additional internal processing sites between mature polypeptide subunits. The dataset columns report the number of proteins and annotated cleavage sites (residue-level), summed across 5 test folds. Precision and recall are reported as mean ± standard deviation across 5 test folds. No benchmark tool was available for chain cleavage prediction. Only proteins with chain annotations in UniProt are included in calculating performance. Cleavage site counts are slightly lower than in **Supplementary Table 2** due to the removal **(Section 2.3)** of a small subset of proteins to enforce the clustering homology threshold.

| Annotation Class | ViralMap | | | Benchmark | | |
|---|---|---|---|---|---|---|
| | PR-AUC | Precision | Recall | Tool | Precision | Recall |
| *Post-translational Modifications* | | | | | | |
| N-Glycosylation | $0.763_{\pm 0.06}$ | $\mathbf{0.701_{\pm 0.06}}$ | $\mathbf{0.844_{\pm 0.07}}$ | (b) | $0.270_{\pm 0.02}$ | $0.827_{\pm 0.01}$ |
| Furin Cleavage | $0.565_{\pm 0.16}$ | $\mathbf{0.506_{\pm 0.13}}$ | $\mathbf{0.690_{\pm 0.31}}$ | (c) | $0.031_{\pm 0.00}$ | $0.440_{\pm 0.04}$ |
| Chain Cleavage | $0.895_{\pm 0.02}$ | $\mathbf{0.918_{\pm 0.01}}$ | $\mathbf{0.859_{\pm 0.03}}$ | — | — | — |
| Disulfide Bond | $0.754_{\pm 0.03}$ | $\mathbf{0.644_{\pm 0.04}}$ | $\mathbf{0.733_{\pm 0.11}}$ | — | — | — |

**Supplementary Table 8.** Baseline classification performance of ViralMap and benchmark tools: **(b)** NetNGlyc **(c)** ProP. This table presents ViralMap's performance using a standard 0.5 probability threshold for all post-translational modification classes (N-glycosylation, furin cleavage, chain cleavage, and disulfide bonds) rather than the F2-optimized thresholds reported in **Table 1**. All metrics are reported as mean ± standard deviation across five test folds. A bold value designates the higher precision or recall metric between ViralMap and the benchmark tool for a given annotation class.

### Supplementary Notes

**Supplementary Note 1: Annotation vector representative selection**

The filtered dataset of 300,406 proteins contained substantial redundancy with certain clusters dominating (e.g., >60,000 entries for HIV-1 envelope protein). Retaining all entries would introduce excessive label noise into model training, while random down-sampling risked removing well-annotated proteins or sparsely annotated proteins containing rare annotation classes. We therefore implemented Algorithm 1 below to select representatives from each cluster, maximizing annotation quality and coverage while minimizing redundancy. To compare quality and coverage across entries, all proteins were represented as fixed-length annotation vectors of 210 dimensions (10 annotation classes × 21 positional bins). This length was determined using the longest protein in the dataset (1,024 residues after the length filter). Using a fixed vector length for all proteins did not have a significant impact on selection since clusters were largely homogenous in length **(Supplementary Fig. 3)**. Each protein's annotation vector was generated by converting all annotations into scores based on their Evidence and Conclusion Ontology (ECO) codes provided in UniProt, then assigning the scores to bins corresponding to the positions of the annotations. Scores were determined based on a five-point scale: experimentally validated annotations received score 4; manual annotations with PubMed references received score 3; manual annotations without PubMed references received score 2; automatic annotations received score 1; and annotations without ECO codes received 0.5. Mapping ECO codes to these annotation scores or tiers is described in **Supplementary Table 1**. Once populated, these annotation vectors had a dual benefit: their magnitudes (L2 norms) captured overall annotation quality, while their directions indicated whether a given protein contained an annotation at a position missing in other representatives. Both norm and direction were used in Algorithm 1 to select representative proteins. Candidate proteins were scored by an 'information gain' metric combining the annotation vector norm and cosine similarity with already selected representatives. The cosine similarity (direction) was used to maximize coverage, since proteins that contributed new information by annotating positions missing in other representatives. This iterative selection continued within each cluster until a hard cap was reached or no remaining protein had annotations that would contribute new information (as measured by a minimum information gain threshold of 1.0, or the equivalent of a single automatic annotation). This reduced the dataset from 300,406 to 8,238 proteins while primarily removing proteins contributing automatic annotations (**Supplementary Fig. 4**). The HIV-1 envelope glycoprotein cluster, for instance, was reduced from 60,348 entries to 17 representatives **(Supplementary Fig. 5)**.

**Algorithm 1**

1: **Input:** Proteins with residue-level annotations across $C = 10$ classes (SP, CC, TM, NG, DR, FR, CH, DB, CY, EX), each with an associated evidence code, and a sequence-identity cluster identifier for each protein.

2: **Construct annotation vectors:**
- ▷ Divide each protein into $B = 21$ bins ($\lceil \frac{1024}{50} \rceil$) of 50 residues each
- ▷ For each protein $p$, initialize annotation vector $\mathbf{v}_p \in \mathbb{R}^{210}$ (10 classes $\times$ 20 bins)
- ▷ For each annotation, increment the corresponding bin(s) by evidence weight $w$: experimental $= 4.0$, manual/PubMed $= 3.0$, manual $= 2.0$, automatic $= 1.0$, unknown $= 0.5$
- ▷ Compute quality score $q_p = \|\mathbf{v}_p\|_2$

3: **Define cluster representative cap:**
- ▷ For cluster of size $n$, set $k_{\max} = \min\big(100, \lceil \log_2(n) \rceil + 1\big)$

4: **Greedy selection within each cluster:**
- ▷ Initialize selected set $S \leftarrow \emptyset$
- ▷ While $|S| < k_{\max}$:
  - For each candidate $p$ not in $S$, compute novelty $\text{nov}_p = \max_{s \in S} \cos(\mathbf{v}_p, \mathbf{v}_s) \quad (0 \text{ if } S = \emptyset)$
  - Compute gain $g_p = \alpha \cdot q_p + \beta \cdot (1 - \text{nov}_p)$ with $\alpha = 1.0$, $\beta = 3.0$
  - Select $p^\star = \arg\max_p g_p$
  - If $g_{p^\star} < \tau = 1.0$, terminate early
  - Otherwise, add $p^\star$ to $S$

5: **Output:** Selected representatives from each cluster, prioritizing proteins with high-quality and non-redundant annotations.
